# “The appetite for freediving differs between Sprague-Dawley and Long Evans rats”

**DOI:** 10.64898/2026.05.04.722625

**Authors:** Laetitia Chambrun, Jorelle Linda Damo Kamda, Laurent Vatrinet, Harquin S. Foyet, Roseline Poirier, Valérie Doyère, Marion Noulhiane

**Author notes:** Present address: Department of Neurosciences, University of Montreal, Montreal, Quebec, Canada (Jorelle Linda Damo Kamda). Equal contribution (Valérie Doyère, Marion Noulhiane). = Co-corresponding authors.

## Abstract

Freediving in rats has emerged as a relevant model to study physiology and neural adaptation underlying submersion mechanisms. However, despite well-established strain-dependent differences in behavior and physiological responses, most studies about freediving rely on Sprague Dawley rats. As the choice of strain could significantly shape experimental results depending on the field of research, we conducted a behavioral comparative study between Long Evans (LE) rats, genetically closer to the Wild Norway rat, with the commonly used Sprague Dawley (SD) strain. We developed an 11-week progressive voluntary freediving protocol involving four distances (from 5 to 11 meters), and assessed the rats’ natural willingness to dive and swim, and identified several parameters for evaluation of their confidence (waiting time before diving, speed), performance capacity (freediving time) and population variability. We found that Long Evans rats were naturally more willing to dive and more confident, compared to Sprague Dawley rats: they showed better performance with longer time underwater and slower diving speed. We also uncover differences in their variability, at trial-to-trial intra-individual and population inter-individual levels, which can guide the choice of one strain over the other, depending on the aim of the scientific inquiry.

**Highlights:**

- LE rats dived more willingly (81% vs 72%) and showed greater confidence.
- LE freedivers waited less, stayed underwater longer, and traveled farther.
- SD rats swam faster and were less confident, with more drop-outs.
- LE rats showed higher variability; SD rats were more consistent.

## 1. Introduction

What makes an animal prone to dive or not? During repeated diving, the brain undergoes multiple exposures to hypoxia. The tolerance to these episodes is variable between species, with aquatic mammals adapted to prolonged submersion, but terrestrial mammals such as rodents and humans having more restricted capacities. Beyond physiological tolerance, diving requires a behavioral decision to suppress breathing and submerge, a choice that varies widely across species and individuals. For example, not every human is willing to train him/herself to reach a record of static apnea lasting more than 10 minutes underwater (World record, Stéphane Mifsud, France 2009) or in dynamic apnea for a distance of 321 meters (World record, Mateusz Malina, Poland 2022). Furthermore, certain populations, such as the Bajau “Sea Nomads” who have marine-dependent lives, have developed, across evolution, the ability to dive during long periods to seek food [1], highlighting inter-individual/population differences even in humans.

Whether diving is a natural behavior as in aquatic mammals and rodents, or a recreational activity as for most humans, it triggers an autonomic reflex common to all mammals. These responses are fairly well described. Historically, the mammalian diving reflex was first described in 1786 by Edmund Goodwyn [2], and later examined in greater detail by Paul Bert [3] in 1870. This field of research was revolutionized by Laurence Irwing [4] in the 1930s and Scholander in the 1940s [5] by characterizing the physiological basis of this reflex. The diving reflex occurs as soon as the head is submerged [6]. It is characterized by several physiological elements such as the activation of both the respiratory chemoreceptors and the anterior ethmoidal nerve [7, 8, 9], leading to a parasympathetic response which is bradycardia, while the sympathetic system triggers the vasoconstriction of vessels thereby redistributing blood flow to the vital organs (i.e heart and brain). The diving reflex was extensively studied in several species [10, 11]: pinnipeds [4], humans [12, 13, 14], rat [15, 16], muskrat [17, 18, 19], whales [20], dolphins [10]. The diving reflex is well-developed in aquatic mammals such as whales and seals, present in semi-aquatic species such as muskrats, but also efficient in terrestrial mammals, including rats and humans.

The foregoing studies initiated the interest in the physiology behind diving and its neural control. An animal model of voluntary repeated hypoxia is also needed for research on long-term modifications, such as brain adaptability, especially as diving as a sport discipline is becoming more popular. In the laboratory, rat models are extensively used, with several strains available. However, the willingness and capacity of rats to be trained to freedive are not documented. The physiological response to and neurobiological mechanisms of water immersion have been studied in the *Sprague Dawley* rat [9, 11, 16, 21, 22], with the Wild Norway rat as an ancestor. In the natural history of the rat, the Norway rat, *Rattus norvegicus,* namely “sewer rat”, is accustomed to living underground and is regularly exposed to water while hunting or moving from one place to another [11, 23], in opposition to the *Rattus rattus*, which lives on rooftops.

McCulloch [21, 22] first developed a voluntary freediving training protocol also using *Sprague Dawley* rats to describe freediving-associated physiological responses such as heart rate, nasal dilation, corticosterone release and brainstem catecholaminergic neurons [11, 15, 16, 22] comparing trained/untrained and forced/voluntary swim/dive conditions. Other studies also used forced immersion to measure arterial pressure (e.g., [24]). As the focus of these studies was on physiological aspects, no measures were reported on freediving abilities in terms of performance, distance, or swimming speed.

However, as the physiological ability for diving may be more or less well-developed depending on the species, one may wonder whether it may also vary at the level of strains or even individuals. As the previous freediving studies have been performed exclusively with *Sprague Dawley* rats [15, 16, 24], no data are available on the potential impact of strain selection. However, strain-dependent differences in physiology and behavior are well documented in other fields of research [25, 26, 27], Thus, the impact of strain selection should be clarified also in freediving studies, especially to take into account which strain might be more willing to engage in and adapted to a freediving task, and thus may exhibit different behavioral performance and stress.

The Wild Norway rat, *Rattus norvegicus,* is the common ancestor to all laboratory rat strains [28]. A study comparing laboratory and wild rats [29] found that wild rat strains dove spontaneously, whereas the three laboratory strains tested (*Sprague Dawley*, *Wistar*, *Brown Norway*) did not. *Long Evans* (LE) rats, also widely used in the laboratory, were created in 1915 by Dr. Long and Dr. Evans by direct crossing of a male wild Norway rat, *Rattus norvegicus,* with a female Wistar [28]. In contrast, *Sprague Dawley* (SD) rats are more distantly related to the Norway rat, as they originate from a cross between a female hybrid Wistar and an unknown hooded male [11, 28]. As LE rats are genetically closer to the wild strain of rats than SDs, they might naturally have a more pronounced interest in water, with more willingness to dive and spend longer times underwater.

The aim of our study was thus to determine whether LE rats might outperform SD rats in freediving, being more willing to accept and more adapted to training, in order to provide a model for further investigation of the neurobiological adaptation to hypoxia related to training in freediving. For this purpose, we created a serpentine watermaze and designed a training protocol inspired by McCulloch with progressive distances from 5 to 11 meters. We analyzed different behavioral parameters to assess the natural inclination for freediving of each strain, their response/confidence when increasing distances and finally their performance and variability between populations. We expected that LE rats could be more comfortable in freediving than SD rats, potentially because of their genetic background.

## 2. MATERIAL AND METHODS

### 2.1 Population

Two different rat strains were used, *Sprague-Dawley* (SD) and *Long Evans* (LE), provided by Envigo and Janvier, respectively. For each strain, 32 male pups (8 days old) were obtained from 5 separate litters (6 to 7 pups per litter). During the pre-weaning period, they were kept in a room dedicated to breeding, with a 12-hour light/dark cycle (8 am to 8 pm) and an average temperature of 21 degrees Celsius. After weaning (21 days or later with a minimum weight > 50g), the rats were distributed into cages of 3 to 4 rats, respecting the litter’s origin, and placed in another colony room at 21°C with a 12h day/night cycle (8 am to 8 pm). The cages were enriched with a red tunnel, sizzle-nest, a cardboard house and a wooden cube; Following weaning, the rats were fed *ad libitum* for 3 days, while we quantified the amount of food eaten. As they grew, the food ration was controlled initially at 95% of the quantity eaten under free access and then adjusted according to their needs. Doses were gradually increased, starting at ∼10g/rat from 2 days after weaning to reach ∼20g each at the end of the experiment when rats were 96 days old (P96). The ration was adjusted in order to observe normal growth and avoid health problems while ensuring that there was nothing left in their cage and not have just eaten at the time of the training session for diving (to avoid diving with a full stomach). Initial weights for the *Long-Evans* rats averaged 50g, reaching a maximum of 330g by the study’s end. Similarly, the *Sprague Dawley* rats began at approximately 60g and concluded at a maximum of 340g. Rats were fed at regular times each day, 30min to 1h after their training.

The experimental protocol described herein conducted at Paris-Saclay Institute of Neuroscience (NeuroPSI) in accordance with the guidelines established by the European Communities Council Directive (2010/63/EU) for compliance and use of laboratory animals and approved by the French Ministry of Research and the French National Committee (2013/6).

### 2.2 Material

The experiment took place from 9am to 5pmIn the experimental room, the light level was set at 20 lux, and the humidity varied between 40% and 55%. To render the task more pleasant for the rats, the water temperature was set between 31°C and 33°C [30], and the room temperature was set at 23°C to maintain the temperature between 22°C and 23°C as the temperature regulation system occasionally fluctuated. Nothing was added in the water.

A serpentine-shaped maze specially designed for this study was created at NeuroPSI (Fig. 1A; supplementary Fig. A.1.) based on McCulloch’s [22] model, to be placed in a 180cm diameter pool, with a water drain. Fully transparent, the maze consists of a starting area, then a path winding around 9 bends, with a lid made of several pieces of Plexiglas preventing the rat from getting out. The lid could be lifted to remove the rat in the event of disorientation or panic. A panic reaction was defined as repeated uncoordinated movements and escape attemps such as scratching at the lid of the maze. Pieces of thick tape at each of the bends, alternating between dotted and continuous tape, intended to help the rat orient itself with its whiskers. There was enough space in the corridors for the rats to turn around (i.e 12cm, Fig. 1). In the starting area, a pedestal allowed the rat to wait “dry” before entering the maze. The exit boxes were removable and could be closed behind the animal if necessary. Several potential exits allow progressive adaptation to apnea and training at increasing distances (D1 = 5m, D2 = 7m, D3 = 9m and D4 = 11m).

**Figure 1:**
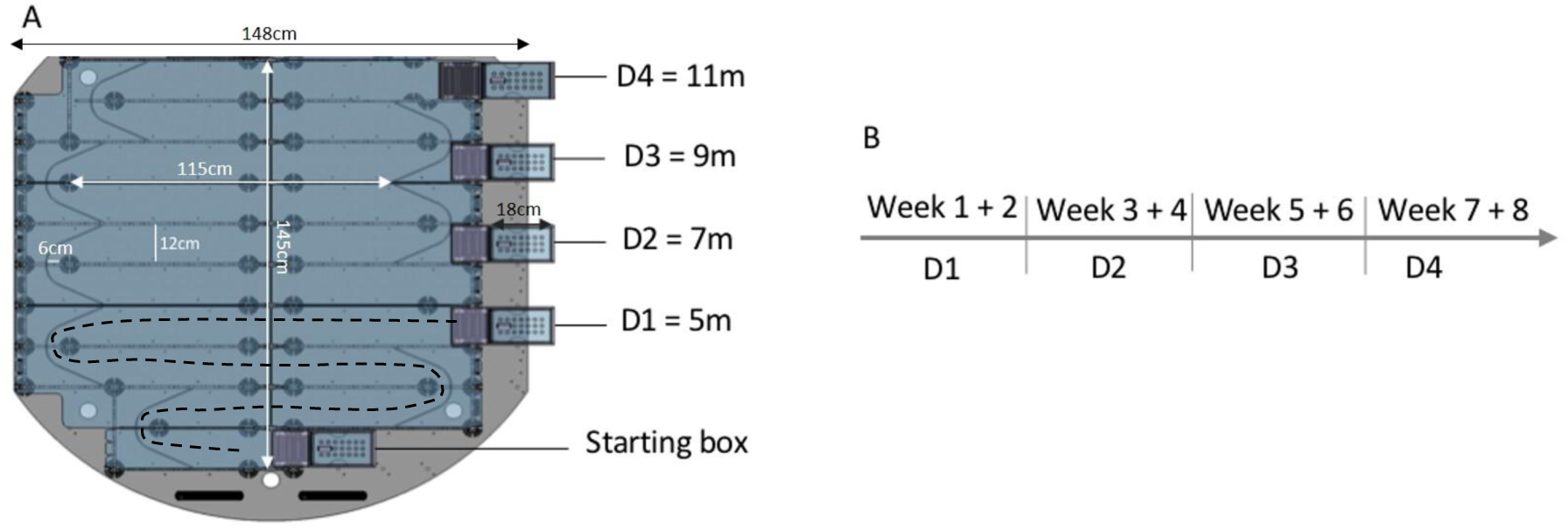
Freediving training in rats. (A): Serpentine-shaped maze layout with distances from the start to each exit. Dot line: example of trajectory at D1 training. (B) Chronology of training distances achieved. (D = distance)

### 2.3 Experimental procedure

#### 2.3.1 Exploration and Pretraining

Two days after weaning, the rats were habituated to being handled and were placed in the serpentine-shaped maze, in groups (cage). Rats had access to the entire maze, with all exits accessible and a reward inside (such as crackers). They spent 10 to 15 minutes in the maze with water 1cm high. To get them out, they were gently guided (by hand) by lifting the plates to an exit box to ensure the pairing of the box with exiting the maze. This exploration phase lasted 3 days.

Then, for 2 weeks, 5 days a week and once per day, the maze was closed beyond distance 1 (D1=5 m), and the rats were placed individually on the pedestal in the starting area. The water level was gradually increased by about 1 cm/day, depending on the rats’ confidence. If the rat did not reach the exit, it was then gently guided to the exit, so that it could be exposed to the complete path. At the end of the 10-day pre-training period, the water reached 10.5 cm, a height that necessitated the rat to swim (instead of walking) to the exit.

To avoid potential stress, we limited the trial duration to a maximum of 5 minutes. If the rat did not leave the pedestal within 180s, it was removed from the starting area and carried to the exit box, and the trial was counted as nogo.

#### 2.3.2 Training

Training sessions progressively increased in difficulty and distance from 5m (D1) to 11m (D4) over 8 weeks, at a rate of 2 weeks per distance (Fig. 1B). As for pre-training, the trial duration was kept to a maximum of 5 minutes (video link: <u>The appetite for freediving differs</u> <u>between Sprague-Dawley and Long Evans rats – Hippomnesis</u>). During training, the reward was given whether the rats performed or not, and can be considered as a treat, to avoid negative association with freediving/swimming and maintain their voluntary participation. The rewards varied according to the difficulty and rat’s taste, from the following list: crackers, corn, coco pops and small pieces of speculoos biscuit Lotus Biscoff^TM^ (Lotus Bakeries, Belgium).

##### (i) Distance 1 (D1 = 5m)

Apnea training began at Distance 1 (5m), for 5 days a week. Each week, the first day was a swimming day (water height h=10.5 cm). From day 2 to day 4, the height of the water was increased above the ceiling lid (h=14 cm), so that the rats had to dive underwater in order to traverse the maze and reach the exit box. If the rat did not dive for 2 to 3 successive days, then the next day, the height of the water was decreased back to the level allowing swimming (h=10.5cm). Then, on day 5 they swam at h=10.5cm. After 2 days of rest, we repeated the same schedule for the second week of training.

At the end of the two weeks of training at D1, different profiles naturally emerged: Freedivers, Swimmers (rats reluctant to dive, but willing to swim) and Nogo (rats reluctant to swim or dive).

##### (ii) Groups redistribution

Before starting the long-distance training (from D2), in perspective of future neurobiological assessments (not reported here), in each strain (32 rats/strain), we formed 3 groups of semi-equal numbers of animals (12 freedivers, 10 swimmers and 10 nogo). While the 12 freedivers remained in their own group, redistribution of some freedivers to Swimmer or Nogo groups, or Swimmers to Nogo group, was done based on the performance during D1 training, and according to the following rules:

– If the rat hesitated one or more times but still attempted apnea, it could remain in the Freediver group or be moved to another group according to the distribution. A rat was moved to the Swimmer group if it did not reach the exit more than once or reach it at least two times but refused to dive at least once. It could also be moved to the Nogo group if the rat reached the exit less than two times and refused/try at least once.
– A swimmer rat was moved to the Nogo group if it exhibited significant reluctance, defined by: hesitation during swimming, failure to reach the exit and inconsistent participation (rats were willing to swim on one session but refused to swim in the next).

##### (iii) Distance D2 (7m) to D4 (11m)

Rats from Swimmer and Freediver groups were trained in the same manner from D2 to D4. Training started with one day of swim (Monday), followed by 3 days of apnea or swim (depending on the group assignment) training and 1 day of swim (Friday). During the transition between each distance, the last swim day was performed at the new distance to acclimate them to it. If, during the training, the rat did not attempt to dive / swim for more than 3 sessions, we considered that the rat gave up and we stopped the training.

Initially, D4 exit was planned on the left-hand side, but this caused a lot of problems, because the rats had gotten into the habit of emerging from the right-hand side of the pool from D1 to D3. The configuration of the maze had thus to be changed during the first session of distance 4, and we added a swimming session to avoid stress related to the spatial reconfiguration As a result, only 4 apnea sessions (common to all rats) could be taken into account.

### 2.4 Analysis

Throughout the training session, we recorded each rat’s passage using a USB 2.0 camera (Stoelting Co., USA; 752×480px, 30fps) placed above the pool, and connected to Anymaze**®** software. In the software we created the apparatus enabling it to track the animal in the maze by delimitating zones: start box, exit box and the serpentine-shaped trace (Supplementary Fig. 1B, 1C).

There was one session per training day, but there may be several attempts until the rat reached the exit, which was then considered as a successful trial (only 1 successful trial per session).

We considered the following measures:

(1) Trial categorization: Trials were categorized as nogo (did not try to dive within 3 min), partially successful (freedive/swim but without reaching the exit within 5 mins) and successful (reach the exit within 5 min). This permitted us to calculate the number of rats in each spontaneously formed category of animals and characterize the freedivers as confident or not-confident during the D1 training:
  – Nogo: the rat did not dive or swim
  – Swimmer: the rat did not dive but accepted to swim
  – Freediver: the rat tried to dive and/or reach the exit at least once over the 6 sessions.
(2) Number of attempts per diving session: the rat could turn around before reaching the exit and return to the pedestal. Several attempts could therefore be carried out in a single session, so we considered the time on the pedestal between each attempt in order to define whether they were one and the same trial, or two different trials. Two attempts were considered as different trials if the rat had spent time on the pedestal between them for more than half of its time in the water.
(3) Waiting time: time spent in the starting box before the first start, on every distance and during swimming day and apnea day. It allowed us to characterize the rat’s confidence.
(4) Speed for the distance traveled, calculated by Anymaze*®* software and the time in apnea or swim for each trial.

#### 2.4.1 Variables

We considered two independent variables: (Strain factor (SF) with two levels (*Sprague Dawley* and *Long Evans*) and Aquatic factor (AF) with 3 levels: Freedivers, Swimmers and Nogo.

Several dependent variables were analyzed:

<u>1 – At Distance D1</u>

(i) Spontaneous Aquatic Behavior (SAB) We analyzed the spontaneous willingness for swimming or freediving during the training sessions at the first distance (D1) and calculated the proportions for LE and SD rats (SF as IV) according to 3 categories: (1) freedivers, rats who were willing to freedive, (2) swimmers, rats who refused freediving but were willing to swim, and (3) nogo rats, those who refused both swimming and freediving. We defined this measure as the proportion of spontaneous aquatic behavior (pSAB). We expected that LE rats would have a higher pSAB freedivers than SD rats.
(ii) Spontaneous Aquatic Confidence (SAC) The spontaneous aquatic confidence (SAC) was evaluated for the freedivers. We considered a rat “confident” if it reached the exit in all 6 freediving sessions during D1 training. A rat was considered “not-confident” if it failed to reach the exit or refused to go at least once during the 6 freediving training sessions. We calculated the proportion of “confident” and “not-confident” freedivers in each strain (pSAC). SD rats might exhibit a higher proportion of not-confident rats compared to LE rats. As another index of confidence, we also analyzed the pre-start latency (waiting time on the pedestal). We expected the waiting time to be longer for not-confident rats.

<u>2 – Preparation to dive/swim when increasing distance</u>

We looked at the preparation to dive/swim (waiting time) across distances from D2 to D4. We had 6 sessions of freediving during D2 and D3, and 4 sessions on D4 (see explanations above).

(i) Waiting time We assessed the extent to which the rats maintained their willingness to dive or swim when the distance increased (from D2 to D4), analyzing the pre-start latency (waiting time on the pedestal) on each successful trial. We assumed that swimmers might maintain a similar waiting time across trials compared to freedivers. We suggested that LE rats (freedivers and swimmers) might have a lower waiting time across trials and distances than SD rats.
(ii) Specificity of diving session We evaluated whether, for the freedivers, the waiting time before the diving session was specific to the preparation to dive, by comparing it with their waiting time on the swimming days before and after each 3-day diving training phase. For comparison, we performed the same analysis for the swimmers. As animals may need some time to prepare for diving, we might expect a higher waiting time at diving sessions compared to swimming sessions, a difference that may increase with difficulty for longer distances.

<u>3 – Freediving behavior at D4</u>

We measured the time spent underwater and distance traveled at D4 (4 sessions) for Freedivers. We also analyzed their speed. No specific hypothesis on strain differences was made.

<u>4 – Population variability</u>

In order to estimate the extent to which the freediving behavior is an individual characteristic of the animal and/or of its strain, we looked at the variability of following parameters – waiting time, freediving time and speed – both between sessions for each animal and between each strain.

(i) Intra-individual variability: we calculated the coefficient of variation (semi-iqr/median) for each animal based on each trial, and then took the median of all the coefficients of variation for each group: CV_intra_ = Semi IQR_rat_ / Median_rat_ = Q3_rat_ – Q1_rat_ / 2*Median_rat_
(ii) Inter-individual variability: we calculated the coefficient of variation (semi-iqr/median) for each group based on each animal’s median: CV_inter_ = Semi IQR_group_ / Median_group_ = Q3_group_ – Q1_group_ / 2*Median_group_
(iii) The intraclass correlation coefficient (ICC) quantifies the proportion of total variance attributable to between-subject (inter-individual) variability. For each group, the ICC was calculated from a linear mixed-effect model including rat identity as a random effect to account for clustering of repeated trials within animals. It was extracted from the variance component of the model at D4. Therefore, a high ICC indicates that variability is mainly explained by inter-individual differences, whereas low ICC indicates a larger contribution of intra-individual variability. To visualize the relative contribution of the intra-individual (within-subject) variability on the inter-individual variability, we calculated the proportion of it based on the ICC: ICC = *σ*(inter)^2^_group_ / *σ*(inter)^2^_group_ + *σ*(intra)^2^_group_

Where *σ*(inter)_group_ corresponds to the variance component between rats and *σ*(intra)_group_ to the variance component within rats.

ICC* = (1-ICC) / CV_inter_

For the waiting time, some animals may have a variability equal to 0. Because of the difficulty of tracking in the starting box, we were not able to have a more precise waiting time when it is close to 0s. This waiting time means that when the rat was put in the starting box, he directly dived/swam.

### 2.5 Statistics

We used the software R Studio (version R-4.5.1.) for each statistical test and ICC. We tested the normality for each database by doing a Shapiro-Wilk test:

– For non-normal distribution we used the following non parametric tests: a Wilcoxon-rank test for non paired variables and a Wilcoxon Mann-Whitney test for paired variables.
– For data with normal distribution, we used a bilateral t-test.

To evaluate the correlation between ‘not-confident/confident’ and the waiting time we calculated the Spearman Coefficient for each strain.

## 3. RESULTS

### 3.1 Ease to dive

#### 3.1.1. Distance 1: Proportion of Spontaneous Aquatic Behavior (SAB) and Confidence (SAC)

##### (i) pSAB

The results indicated that there was a tendency to have more LE Freedivers compared to SD rats (81% *vs.* 72%; Fig. 2A). For the non-diving rats, SD rats were either swimmers (12%) or nogo (16%), whereas LE rats refusing to dive also refused to swim (19% nogo). Thus, the main difference between strains was the fact that we found only two profiles in LE rats (freedivers and nogo), in contrast to the three different profiles in SD rats (freedivers, nogo and swimmers; Fig. 2A).

**Figure 2:**
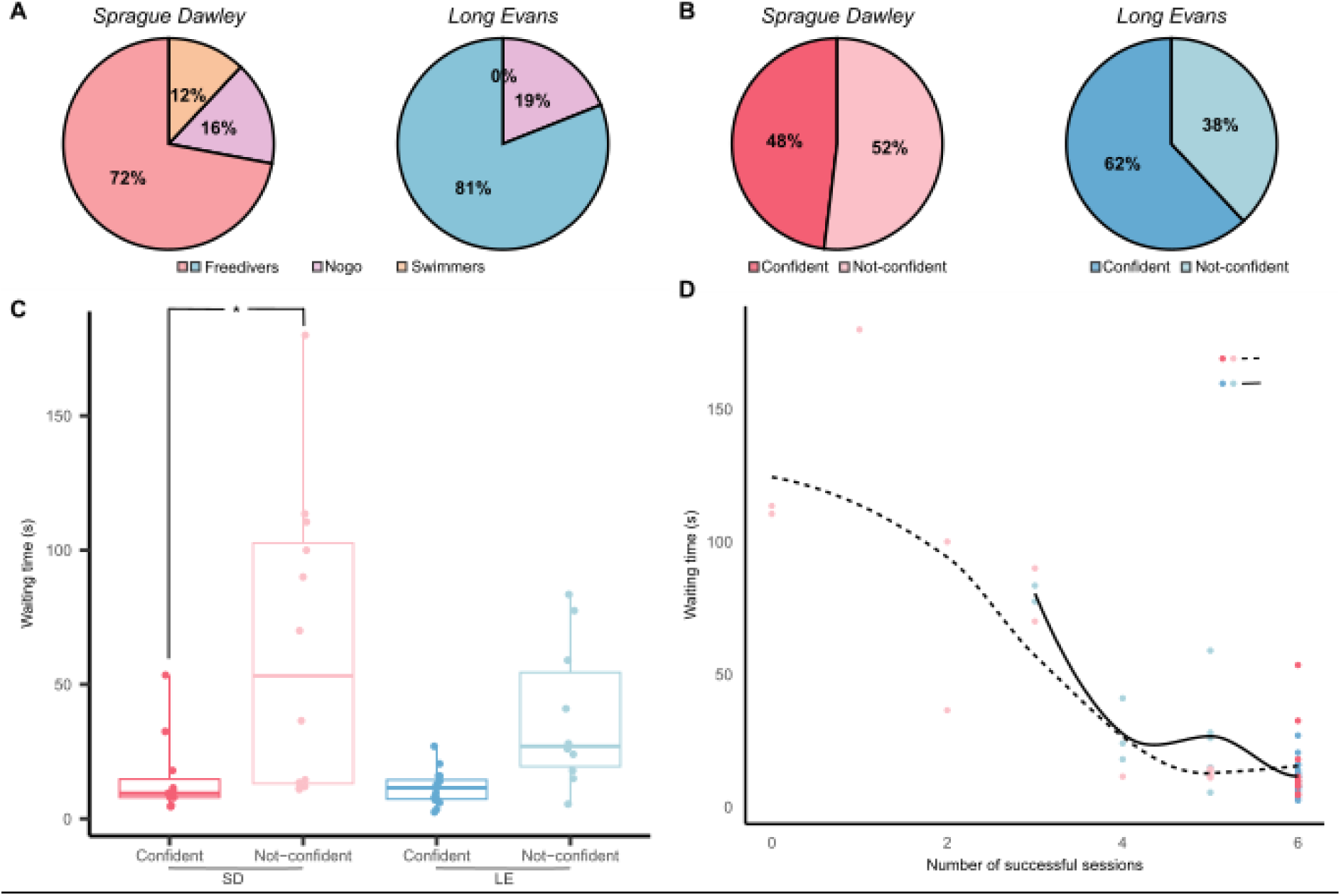
Characterization of the rats’ natural appetite for freediving at distance D1. (A) Natural proportion of freedivers, swimmers, nogo for each strain after D1, from 32 *Sprague Dawley* (SD) rats and 32 *Long Evans* (LE) rats. (B) Proportion of confident and not-confident freedivers rats for each strain during D1. (C) Waiting time (s) (boxplots showing median, interquartile, min and max) before first start for confident and not-confident LE and SD Freedivers. (D) Spearman correlation plot between the median of the waiting time (s) and the number of successful sessions during D1 of each rat (confident and not-confident) for each strain.

##### (ii) pSAC

Based on the observed unwillingness to freedive successfully at least once during training at distance 1, the results showed that among the 26 LE Freedivers, there were more rats classified as “confident” (16 rats, 62%) than not-confident (10 rats, 38%; Fig. 2B). In contrast, there were 11 confident and 12 not-confident rats among the 23 SD Freedivers, representing 48% and 52% respectively.

The amount of time the rats spent in the starting area (on pedestal), or waiting time, can also be representative of the rat’s hesitation (Fig. 2C). In both LE and SD freediving rats, we observed that the confident rats spent less time in the start zone before diving than the not-confident rats, with the difference reaching significance in the SD (W=17.5, P =0.003) and in the LE (W=22, P=0.002). Not-confident LE rats tended to spend less time on the pedestal than not-confident SD rats, although the difference did not reach significance (W = 45.5, P=0.3557).

To assess whether the waiting time could be related to some level of confidence, we analyzed the correlation between the waiting time and the number of successful sessions taking into account all freedivers (Fig. 2D). We obtained for LE and SD a coefficient of correlation of ρ_LE_ = – 0.6587 (P < 0.001) and ρ_SD_ = –0.7726 (P < 0.001)(Fig. 2D), respectively, showing that, in both groups, the lower the waiting time, the more likely the animal will dive successfully for the majority, if not all, of training sessions. When focusing only on the not-confident animals (n_LE_=10 LE, and n_SD_= 12 SD), a strong correlation was observed for SD rats ρ_SD not-conf_ = – 0.87 (P < 0.001), whereas the correlation in LE rats was weaker and not statistically different (ρ_LE not-conf_ = –0.55 (P =0.09), likely due to the higher number of successful diving sessions in LE rats. These results confirm that SD rats were less confident as reflected in their performance during D1.

##### (iii) Drop-outs

None of the LE rats, swimmers or freedivers, gave up when increasing the distance during training. In contrast, one SD swimmer gave up and became Nogo during the training at D3. Also, at the longest distance (D4), three SD Freedivers gave up and became Swimmers, and two SD swimmers became Nogo, after more than 3 unsuccessful sessions. We eliminated those rats for the following analysis (D2 to D4), leaving 12 LE and 9 SD Freedivers and 10 LE and 7 SD swimmers.

#### 3.1.2. Willingness to freediving when increasing maze length: waiting time

During the training, and for each distance, days 1, 5 and 9 are swimming days for both groups (Freediver and Swimmer), while day 2-4 and 10-12 are freediving/swimming days according to the groups. We characterized the willingness when increasing maze length and compared it to swim days.

##### (i) Waiting time (D2 to D4)

From D2 to D4, neither SD nor LE swimmers significantly changed their waiting time on the pedestal before the first start as the difficulty increased (D2 vs. D3: V=1, P=0.10 and V=1.5, P=0.07; D3 vs. D4: V=9, P=0.83 and V=5.5, P=0.17; for SD and LE, respectively, Fig. 3A). When comparing the two strains, SD swimmers tended to wait more on the pedestal than LE swimmers at each distance, although none of the differences reached significance (SD vs. LE at D2: W=16.5, P=0.07 / D3: W=17, P=0.08 / D4: W=17.5, P=0.09; Fig. 3A).

**Figure 3:**
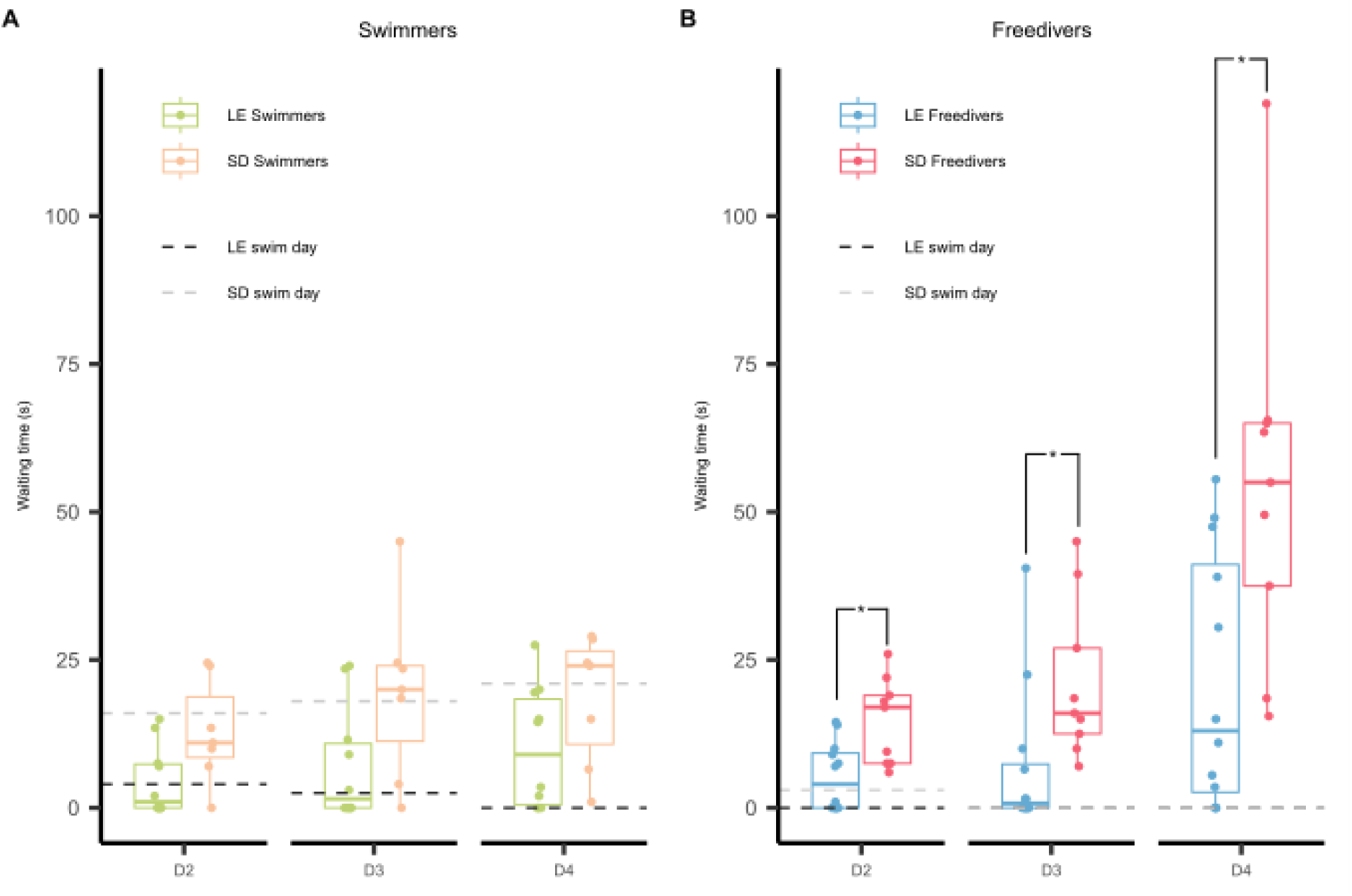
Evolution of the rat’s willingness to dive when increasing the distance. **(A)** Boxplot showing median of waiting time (s), quartiles, min and max for the rats of the Swimmer groups (days 2 to 4 of each week). For comparison with freedivers, horizontal dashed lines show median waiting time (s) during swimming sessions common to all groups (days 1, 5 and 9 of each distance). (B) Boxplot showing median of waiting time (s), quartiles, min and max for the rats of the Freedivers groups during freediving sessions (days 2-4 of each week). Horizontal dashed lines show median waiting time during swimming sessions (days 1 and 5 of each week).

In contrast, both SD and LE Freedivers increased their waiting time before the first start with lengthening distances. When the distance increased, SD, but not LE, rats tended to increase their waiting time when transitioning from D2 to D3 (V=7.5, P=0.08 and V=15, P=0.72, for SD and LE, respectively; Fig. 3B), while both strains changed their waiting time when the distance increased from D3 to D4 (D3 vs. D4: V=0, P<0.01; V=1, P<0.01 for SD and LE, respectively). In addition, SD Freedivers were waiting significantly more than the LE Freedivers at each distance (SD vs. LE at D2: W=18, P=0.011 / D3: W=15.5, P<0.01 / D4: W=16, P<0.01; Fig. 3B).

Thus, SD rats globally waited longer on the pedestal than LE rats, especially the freedivers, suggesting a lower confidence and/or motivation. With increasing difficulty (i.e. distance), both freedivers strains were impacted, but sooner for SD (from D3) than for LE (from D4).

##### (ii) Specificity of preparation to dive

To evaluate whether the increased waiting time was specific to the diving session, we looked at the swimming days before and after each 3-day diving training phase. Importantly, as the last swimming session of the previous distance was at the new distance, these results are independent of the novelty effect.

SD Freedivers were spending much less time on the pedestal on swimming days compared to freediving days at each distance (D2: V= 43, P =0.01; D3: V = 45, P < 0.01; D4: V= 45, P < 0.01). In contrast, LE Freedivers waited less during swimming days compared to diving days only at D4 (D2: V =23, P=0.15; D3: V = 18, P=0.14; D4: V=45, P<0.01). Both strains had a similar waiting time during swimming days (SD vs. LE, at D2: W = 33.5, P =0.11; D3: W=48.5, P=0.65; D4: W= 48, P=0.62; Fig. 3B), without changing across distance (SD, D2 vs D3: V=17, P= 0.2; D3 vs D4: V=1, P=0.42 / LE D2 vs. D3: V = 2, P=0.78; D3 vs D4: W=1, P=0.42; Fig. 3B).

These results show that the differences between strains are mainly linked to freediving, as LE and SD swimmers were not statistically different.

#### 3.1.3. Freediving behavior at distance D4

We characterized the freediving behavior during the final 11-metters distance (D4) of the training.

##### (i) Speed

SD Freedivers were significantly faster than LE Freedivers (SD vs. LE at D4: W=24, P=0.03; Fig. 4A), suggesting a higher motivation to exit at the end of the maze. For both strains, Swimmers had a significantly lower speed, due to the differential air/water resistance (Freedivers vs. Swimmers, SD M_freedivers_=0.41 vs M_swimmers_=0.23, W=63, P<0.01 / LE M_freedivers_=0.38 vs M_swimmers_=0.22, W=120, P<0.01), without significant difference between strains (W=34, P=1).

**Figure 4:**
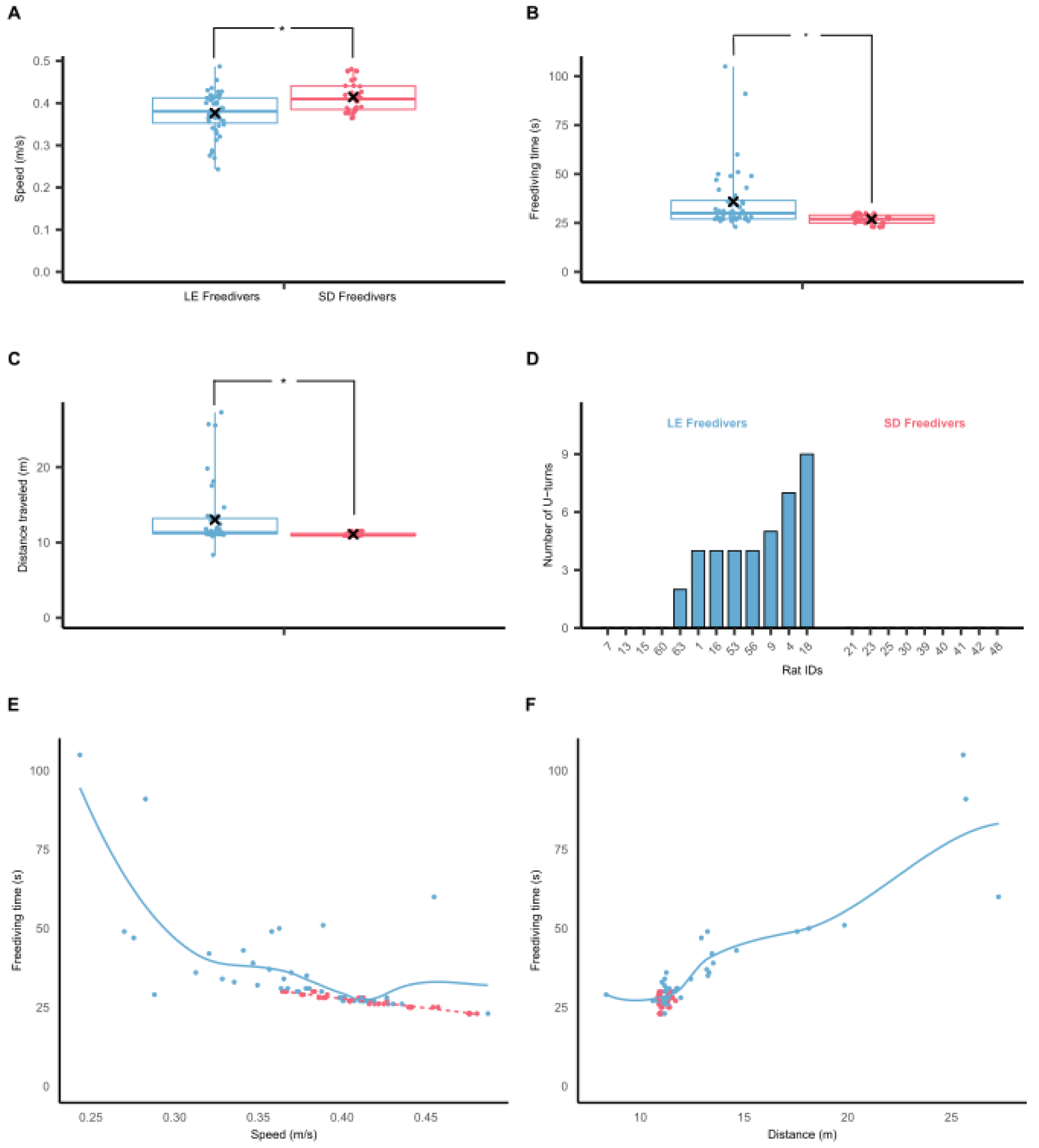
Freediving during each successful trial at D4. **(A)** Boxplot showing the evolution of median speed (m/s), quartiles, min and max for Freedivers. **(B)** Boxplot showing median, quartiles, min, max and mean (cross) of the time spent in apnea (s). Each point corresponds to a trial in a successful session from one rat. (C) Boxplot showing the evolution of median distance traveled (m), quartiles, min and max for Freedivers. (D) Histogram showing the total number of U-turns made at D4 by each individual rats (IDs). **(E)** Spearman correlation plots between the freediving (time) and the speed for each strain. Each point corresponds to a trial in a successful session from one rat. (F) Spearman correlation plots between the freediving (time) and the distance traveled for each strain. Each point corresponds to a trial in a successful session from one rat.

##### (ii) Time spent in apnea

LE Freedivers spent significantly more time in apnea than SD Freedivers (t = 3.37, df=81, P<0.01; Fig. 4B). Interestingly, there was a strong disparity between the two groups, with values clustering around 25 seconds for SD Freedivers, while LE Freedivers diving time ranged from 20 seconds to more than 100 seconds.

##### (iii) Distance traveled

Even though the maze has a fixed length, the distance traveled may be variable, as animals are free to make U-turns as much as they want. LE Freedivers significantly traveled more than SD Freedivers (W=1346.5, P<0.001, Fig. 4C), with a strong disparity for LE Freedivers compared to SD Freedivers. We observed during the entire training that LE rats tended to make more U-turns than SD rats. At distance 4, there was a strong disparity between the two strains. Some SD rats stopped making U-turns, while LE rats continued, with some making frequent U-turns mostly in several sessions (number of u-turns > 2) and others none (Fig. 4D).

##### (iv) Relationship between speed, freediving time and distance traveled

We then investigated the relationship between speed, distance traveled and diving time, through the calculation of the Spearman correlation coefficient.

There was a strong negative correlation between the time underwater and the speed (Fig. 4E) for both SD Freedivers (ρ_SD Freedivers_=-0.95, P<0.001), and LE Freedivers (ρ_LE Freedivers_=-0.77, P <0.001). Thus, the faster the animal was, the shorter the diving time, an intuitively logical relationship. More interestingly, the time spent in apnea by LE Freedivers was strongly modulated by their distance traveled (ρ_LE Freedivers_=0.75, P <0.001, Fig. 4F), whereas there was no such correlation for SD Freedivers (ρ_SD Freedivers_=-0.03, P=0.82, Fig. 4F). With regard to the relationship between the distance traveled and speed, there was a weak but significant negative correlation for LE Freedivers (ρ_LE Freedivers_=-0.32, P =0.02) whereas no significant correlation was found for SD Freedivers (ρ_SD Freedivers_=0.31, P =0.06). Thus, LE rats that traveled longer distances, depending on the number of U-turns which primarily affects the distance traveled, tended to swim slightly more slowly.

The higher variability observed in LE rats led us to investigate the source of variability (intra and inter) in each strain for each studied parameter at D4.

### 3.2 Freedivers’ variability at distance D4

#### (i) Intra-individual variability

The source of variability in freediving behavior may stem from differences from one trial to another for each individual rat. We compared the two strains for their consistent behavior during the four trials at distance D4 by calculating for each parameter (waiting time, freediving time, speed, distance traveled) the intra-individual coefficient of variation (CV_intra_, Fig 5, left panels).

**Figure 5:**
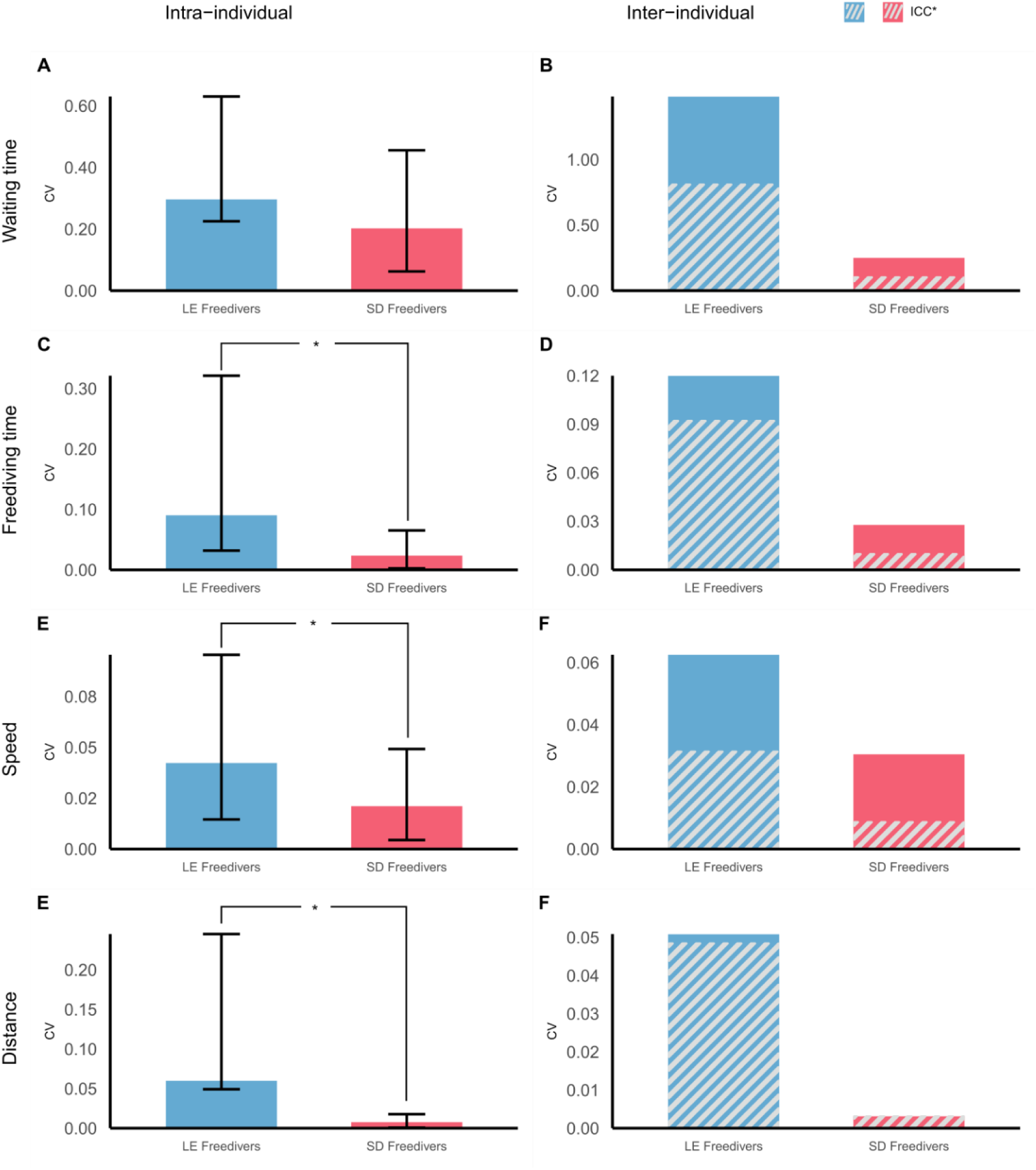
Strain variability at D4 for three behavioral parameters. Left panels: Barplots showing median intra-individual coefficient of variation with quartiles for LE and SD Freedivers at D4, for waiting time (A), freediving time (C), speed (E) and distance traveled (G). Right panels: Barplots showing median inter-individual coefficient of variation for LE and SD Freedivers at D4 for waiting time (B), freediving time (D), speed (F) and distance traveled (H). Stripes represent the relative contribution the intra-individual variability brings to the intervariability CV, calculated through the intraclass correlation coefficient *(ICC\**_intra_ *= (1-ICC) x CV*_inter_).

While LE and SD Freedivers were consistent in their waiting time across trials (W=66, P=0.41, Fig. 5A), they differed significantly in freediving time (W=93, P<0.01, Fig. 5C), speed (W=86, P=0.02, Fig. 5E) and distance traveled (W=89, P=0.02, Fig. 5G), with LE rats showing higher variability than SDs.

#### (ii) Inter-individual variability

For the four behavioral parameters (Fig. 5, right panels), LE Freedivers showed a higher inter-individual coefficient of variation than SD rats. However, the difference in inter-individual variability could in part be explained by the rat’s variability across trials. For the waiting time (Fig. 5B), approximately half of the inter-individual variability came from the intra-individual variability for both strains (ICC_inter_ LE=0.45 and ICC_inter_ SD=0.59). For the freediving time (Fig. 5D), LE Freedivers variability was mainly explained by the rat’s variability from one trial to another (ICC_inter_ LE=0.23), whereas SD Freedivers intra-individual variability represented a small part of group’s variability (ICC_inter_ SD=0.65). For speed (Fig. 5F), 41% of the inter-individual variability was explained by the intra-individual variability in LE Freedivers (ICC_inter_ LE=0.59), whereas it was mainly reflecting inter-individual variability in SD Freedivers (ICC_inter_ SD=0.71). The inter-individual variability in the distance traveled is totally explained by the intra-individual variability for SD Freedivers (ICC_inter_ SD=0) and at 96% for LE Freedivers (ICC_inter_ LE=0.04).

These results suggest that LE Freedivers were not consistent in their own behavior from one trial to another (especially for the freediving time and distance traveled) and were a more heterogeneous population than SD Freedivers.

## 4. DISCUSSION

The aim of the present study was to assess whether rat’s strain may have a significant impact when studying voluntary freediving in preclinical models. Designing a new protocol, inspired from McCulloch’s 5m-maze [22], we succeeded in training both strains of rats to voluntary freedive repeatedly up to 11m. Freediving abilities of rats from two strains – *Long Evans* (LE) and *Sprague Dawley* (SD) – were compared by analyzing several parameters across freediving distances (i.e., waiting time before diving, speed and time in apnea). We found consistent evidence that LE rats were the most suitable strain for freediving: they exhibited shorter waiting time before diving and spent more time underwater, compared to SD rats. Throughout training, LE rats exhibited a greater confidence and ease in the water (with lower speed and more distance traveled, indicating no rush for exiting the water). However, LE rats appeared to be more heterogeneous, both within their own behavior from session to session and across individuals. Their greater inter-individual variability was mostly explained by their higher trial-to-trial variability.

### 4.1 Genetics differences

According to literature, the diving reflexes can be heritable, such as bradychardia [31], suggesting that genetic background may contribute to the strain-differences observed. LE rats and SD rats have different genetic backgrounds, due to the difference in male breeders, a wild grey rat, *Rattus Norvegicus,* for the LE strain, and an unknown hooded rat for the SD strain. Interestingly, in the Stryjek’ study [29], the authors compared the behavior of wild type strain of rats and 3 laboratory strains (Wistar, Brown Norway and SDs rats), when they were housed in one tank connected to a water-filled tank during 72 hours. Among the 12 SD rats, none of them swam or dived, demonstrating a lack of interest for aquatic exploration, while 16% of the wild strains’ rats swam and dived. These results are consistent with the idea that genetic background may play a role in the differences in spontaneous aquatic behavior.

These differences may also be interpreted in terms of evolutionary trade-off. Traits associated with increased exploration (including diving) and adaptation to challenging environment may provide advantages in certain ecological contexts [32, 33]. For example, in wild animals, diving could be advantageous for resource acquisition/variety and in consequence their survival [34, 35, 36]. However, reduced exploration and greater avoidance may represent an alternative strategy by limiting threats and unnecessary energy consumption [36, 37]. *Long Evans* and *Sprague Dawley* strains may have had divergent selection histories, with LE rats potentially conserving behavioral characteristics in favor of exploration and environmental adaptability, and SD rats displaying a more conservative strategy, potentially linked to their albinism phenotype leading to diminished vision and by consequence less exploratory behavior (see below).

However, despite their differences in ancestors and possible evolutionary trade-off, LE and SD rats have undergone many generations of laboratory breeding, rendering difficult to determine to what extend these differences reflect an ancestral characteristic, strain-specific genetic background or other consequences of domestication.

Differences in genetic backgrounds may also lead to variability within strains. Several studies showed that there is genetic and phenotypic variability between different suppliers in *Sprague Dawley* rats [39, 40]. In particular, Gileta *et al*. [40] demonstrated significant genetic differences not only between suppliers but also among sub-groups from the same supplier.

Our rats came from different suppliers, with *Long Evans* obtained from Janvier Labs and *Sprague Dawleys* from Envigo. Thus, in our study, we cannot exclude that the origin of the animals (supplier) and/or their litters may have influenced the results. Additionally, potential differences in reproductive traits, such as litter size between both strains, could contribute to the variability within and across strains [41], although we deliberately asked the suppliers to provide litters of approximately the same size (6-7 male pups per dam). Unfortunately, we do not know whether the litters were left intact or subject to reshuffle by the suppliers (cross-fostering, merging of litters) before their delivery. However, in our lab, we have repeatedly observed higher rates of drop-outs and lower motivation for freediving in SD rats compared to LE rats (unpublished data). Furthermore, difference in intra-strain variability has also been reported previously, even with strains coming from the same supplier [42], with higher variability in pre-pulse inhibition in LE rats, compared to SD rats. These observations concur with our results, pointing to a behavioral difference between LE and SD strains.

Strain-specific genetic diversity could therefore underlie behavioral flexibility and variability observed during freediving, leading to greater confidence and adaptability of *Long Evans* rats compared to *Sprague Dawley* rats. Nevertheless, alternative explanations such as phenotypic differences and individual behavioral strategies should be considered.

### 4.2 Phenotypic differences: visual acuity

Genetic diversity between strains is reflected by phenotypic differences, which can be observed through visual perception or locomotion.

We know that rats rely heavily on their sensory systems, and visual acuity could have influenced how each strain anticipates and navigates in the maze. We noticed that rats from both strains kept their eyes open during freediving, suggesting that they attempt to use visual information, even if its effectiveness in such conditions is uncertain. SD rats have a lack of pigmentation because of a genetic mutation, causing albinism, in contrast to LE rats. This difference may play a role in the observed results. For instance, Prusky *et al.* [43] tested the visual acuity between pigmented and un-pigmented rats including SD and LE rats on a visual water task. It was a visual discrimination test in which the rat swims in a trapezoid-shaped pool toward one of two screens, one displaying a grating pattern and the other blank, and learns to choose the patterned screen to reach a hidden escape platform beneath the water. They showed a main difference in visual acuity: LE rats had approximately 50% better visual acuity than SD rats [43]. In addition, it has been demonstrated that LE rats were more accurate in locating the hidden platform in the classical spatial memory task using environmental cues (Morris Water Maze task) than SDs [44], a difference that may stem from the visual system. SD rats also showed a higher sensitivity to light intensity/change and brightness [45]. We cannot exclude the possibility that the small reflection of light in the water may have disturbed their visual acuity, with higher impact on already visually impaired SDs. It could be as if the animals were going to dive blindly, leading to reduced confidence and longer waiting time. Thus, it is conceivable that the differences in visual acuity may have impacted their ability to correctly distinguish the maze walls, turns, or even the exit, and, consequently, their confidence in diving.

### 4.3 Phenotypic differences: locomotion and exploratory behavior

On the other hand, the motor strategy employed may have had a strong impact on several parameters (i.e freediving time, distance traveled, speed) and several studies have shown that the two strains have different exploratory behavior [42, 44]. Turner *et al* [42] performed several tests (hole board, radial arm maze, elevated plus maze) which revealed greater locomotion in LE rats, especially when looking at the distance traveled in each test, compared to SD rats. Some studies demonstrated a difference in motor control depending on strains [46, 47, 48]. For example in reaching strategies [46], it was described that SD rats showed more rigid and restrictive quality of movement than LE rats to grab a pellet on a platform. The authors also suggested that SD rats did not present completed extensor/flexor movements, contrary to LE rats.

Better motor skills and/or higher exploratory inclination may have contributed to the fact that LE rats spent more time underwater, diving at a lower speed and traveling more, with a high variability among animals and sessions, in part related to the number of U-turns the rat made in our serpentine-maze. A speed heat-map analysis performed on some animals making systematic U-turns (not shown) indicated that there was no abnormal change in speed before a U-turn, suggesting that there was no anticipation for U-turn, and thus no systematic strategy in the diving route. In fact, we observed a direct link between the time in apnea and the distance traveled for LE Freedivers. This was not the case in SD Freedivers, which were on the contrary systematic in their diving route, meaning that they were “rushing” to the exit, with a low intra– and inter-individual variability on the distance traveled. The fact that LE rats made more U-turns, but in an unpredictable way, might be related to their more precise and less rigid movement strategies. Therefore, it is plausible that LE rats feel more comfortable in water, as their swimming strategy may be better adapted than that of SD rats, leading to a higher variability in their behavior, with U-turns related to impulse or desire of the moment.

### 4.4 Individual differences

An important finding in the present study is the difference in inter-individual variability observed within both strains, particularly in *Long Evans* rats. Such variability reflects more individualities in strategies adopted when facing an underwater, rather than strain characteristics only. In the study of Krafft [49], when Wistar rats had to dive to get food pellets on the other side of the tank, two profiles emerged, with carrier-rats providing food to non-carrier rats. In addition to the different social profiles, individuals were showing different strategies. For example, when losing a pellet into water, some rats were diving to retrieve it, while others were simply returning to get another pellet or even sometimes abandoned the attempt. The non-carriers were also showing different behavior, some sharing the food, while others stealing it from carrier. These observations suggest that underwater behavior is shaped not only by strain-related characteristics, but also by individual differences in behavior.

Individual behavioral variability has been described in several studies. One interpretative framework of the behavioral variability in our study relates to the distinction of goal-tracking and sign-tracking phenotypes. In Pavlovian conditioning paradigms, goal-trackers approach the location of the reward whereas sign-trackers focuses on the stimulus predicting the reward. Studies conducted in both strains (SD and LE) showed a broad distribution of both phenotypes, influenced by strains characteristics and animal suppliers [40, 50, 51]. While *Long Evans* rats show a higher proportion of sign-trackers [52, 53], the behavioral profile of *Sprague Dawley* rats appears to depend strongly on the supplier. For example, Charles River SD rats predominantly display goal-tracking behavior, while Harlan SD rats are more frequently classified as sign-trackers [50]. In addition, Srey et al. showed in *Long Evans* rats that extended training can promote a shift toward a sign-tracking behavioral phenotype [54]. Interestingly, LE rats show greater impulsivity and addictive profile compared to SD rats [55]. Impulsive/addictive behavior has been associated with sign-tracking with different interindividual profiles [54, 56].

Although our paradigm differs fundamentally from classical Pavlovian conditioning tasks, the behavioral variability observed here may reflect these goal– and sign-tracking tendencies. In our paradigm, the goal would be the exit box leading to the obtaining reward, whereas the maze itself may provide salient environmental stimuli. From this perspective, the higher number of U-turns performed by *Long Evans* freedivers at Distance 4 may reflect increased exploration of the maze environment, a sign-tracking-like behavioral profile. In contrast, the absence of U-turns in *Sprague Dawley* freedivers suggests a more direct, goal-oriented strategy.

Overall, the inter-individual variability observed in our paradigm may potentially be related to differences in sign– and goal-tracking profiles, although this remains to be tested. Nevertheless, the high intra-individual variability observed in LE rats can hardly be explained by these behavioral analogies.

### 4.5 Forced vs. voluntary

Studies using forced immersion, such as Panneton *et al.* [16], showed that SD rats lose consciousness within ∼70 seconds of immersion and need ∼25 seconds to start breathing once removed. In voluntary diving, however, rats remained conscious and resumed breathing immediately, highlighting the critical role of voluntary participation during underwater tasks. However, to our knowledge, there is no report in the literature regarding effect of immersion forced test in LE rats. The difference between voluntary and forced conditions might relate in part to the difference in the level of stress or anxiety triggered by these two conditions. As, in our experiment, LE rats were capable of diving for as long as 105 sec, whereas the maximum in SD rats was 30 sec, one may wonder whether it reflects different stress or anxiety levels between the two strains. Sanchis-ollé *et al.* [57] demonstrated that *Long Evans* rats seemed to have a higher response to stressors, with a high production of ACTH during forced swim test.

In addition, Snyder *et al.* [58] showed that *Long Evans* were more responsive to chronic intermittent hypoxia with an association between oxidative stress and HPA hyperactivity. However, during elevated plus maze tests and light-dark emergence tests, which evaluate the rat’s anxiety, they presented a reduced anxiety by spending more time in the open section than did SD rats [42]. Conversely, another study [59], using an elevated plus maze test, highlighted the absence of difference in anxiety in this task between the two strains. In our voluntary freediving task, LE rats spent more time underwater, with U-turns related to their momentary state, and were slower than SD rats, which may reflect greater comfort and reduced stress/anxiety when free to choose.

### 4.6 Conclusion

Overall, our findings highlight the importance of considering genetic and phenotypic characteristics when interpreting behavioral and physiological responses during complex tasks such as voluntary freediving. Future studies comparing inbred and outbred strains, as well as pigmented and non-pigmented rats, would help disentangle the respective contributions of vision, stress physiology, and motor ability.

Our study points to the importance of taking into account both inter– and intra-strain variability in behavior to have a better representation of a population, for making an informed choice depending on the research field. Because *Long Evans* rats demonstrated a high inter– and intra-individual variability, they might be more appropriate than *Sprague Dawley* rats for studying traits with strong natural variation (e.g. predisposition to voluntary freediving). Conversely, to study physiological parameters (e.g mammalian diving reflex), *Sprague Dawley* rats should be privileged because of their lower variability allowing more stable physiological analysis. Importantly, such variability is not restricted to rodent/animal models but it can also be observed in human populations. The Bajau, called “Sea Nomads”, who need to dive for their survival (food), have developed physiological adaptations (spleen size, oxygen storage), highlighting the relevance of considering natural diversity when interpreting experimental findings.

However, it is also important to acknowledge current limitations in physiological data. While previous work [7,15] has provided physiological data concerning the response to apnea in SD rats, comparable knowledge remains non-existent for LE rats. This gap restricts direct physiological comparisons and underscores the need for future studies integrating both behavioral and physiological approaches across strains. Given our findings, we believe that LE rats may be the primary choice for investigating the brain adaptability and neurophysiology underlying voluntary freediving as an extreme sport.

## Supplementary Figure

**Figure A.1:**
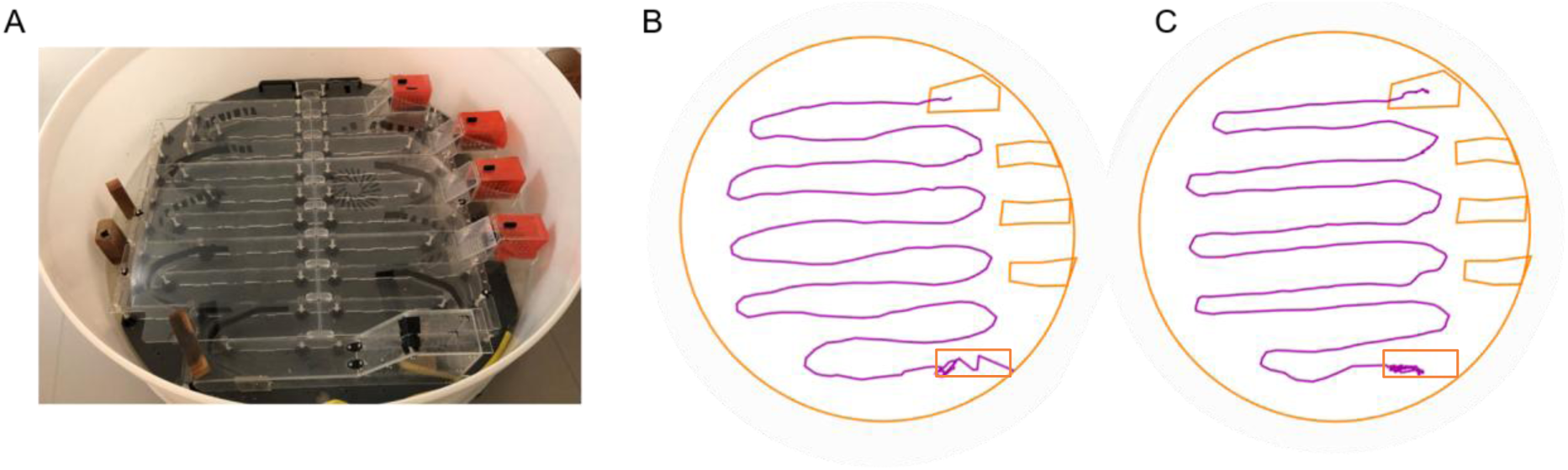
(A) Photography of the serpentine-shaped maze (red = exit boxes). (B) Tracking of a *Long Evans* rat during diving at D4 from the starting box to the exit box. (C) Tracking of a *Sprague Dawley* rat during diving at D4 from the starting box to the exit box.

## Authors Contributions (CReDit)

Laetitia Chambrun: Conceptualization, formal analysis, investigations, methodology, visualization, writing (original draft)

Jorelle Linda Damo Kamda: Conceptualization, methodology

Laurent Vatrinet: Conceptualization, Ressources

Harquin S. Foyet: Funding acquisition, Supervision *(of Linda)*

Roseline Poirier: Conceptualization, methodology, Writing (review & editing)

Valérie Doyère: Conceptualization, funding acquisition, methodology, project administration, supervision, writing (original drafts), writing (review & editing)

Marion Noulhiane: Conceptualization, Funding acquisition, methodology, Project administration, supervision, writing (original drafts), writing (review and editing)

### Fundings

Recurrent funding from CNRS (VD), Paris-Saclay University, INSERM, CEA (MN); Bourse Eiffel Excellence, Paris-Saclay University (France) and DSCA (Dispositif de Soutien aux Collaboration avec l’Afrique du Sud), CNRS (France) (VD, HF); PhD Grant B2V des mémoires (LC, VD, MN)

## Acknowledgement

We thank the NeuroPSI’s behavioural platform, PSI-CO, for providing access to the experimental facilities and equipment. We also thank the NeuroPSI animal caretakers for their daily engagement during the experiments.

