## Supplementary Figure 1 for "“The appetite for freediving differs between Sprague-Dawley and Long Evans rats”"

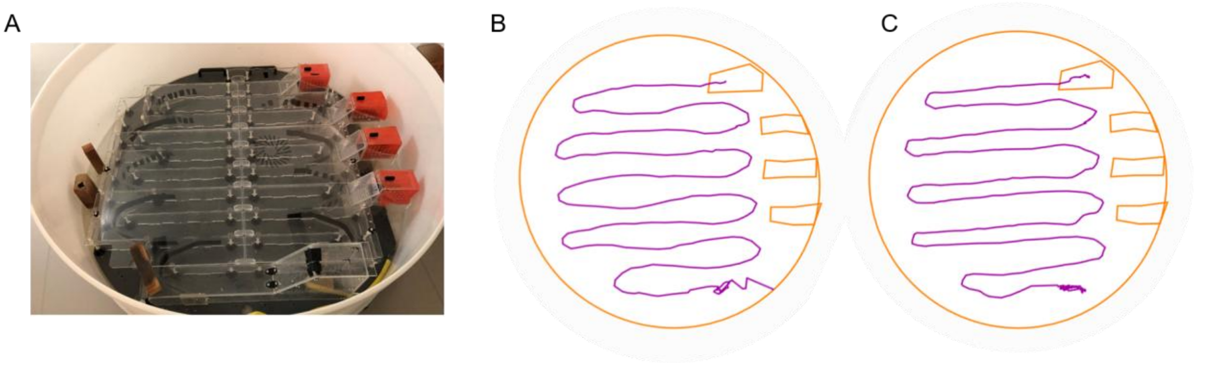


**Supplementary Figure 1:** (A) Photography of the serpentine-shaped maze (red = exit boxes). (B) Tracking of a *Long Evans* rat during diving at D4 from the starting box to the exit box. (C) Tracking of a *Sprague Dawley* rat during diving at D4 from the starting box to the exit box.
